# 7β-hydroxysteroid dehydratase Hsh3 eliminates the 7-hydroxy group of the bile salt ursodeoxycholate during degradation by *Sphingobium* sp. strain Chol11 and other *Sphingomonadaceae*

**DOI:** 10.1101/2025.01.23.634595

**Authors:** Phil Richtsmeier, Ruslan Nedielkov, Malte Haring, Onur Yücel, Lea Elsner, Rebekka Herdis Lülf, Lars Wöhlbrand, Ralf Rabus, Heiko Moeller, Bodo Philipp, Franziska Maria Mueller

## Abstract

Bile salts are steroids with distinctive hydroxylation patterns that are produced and excreted by vertebrates. In contrast to common human bile salts, ursodeoxycholate (UDCA) has a 7-hydroxy group in β-configuration and is used in large amounts for treatment of diverse gastrointestinal diseases. We isolated the 7β-hydroxysteroid dehydratase Hsh3 that is involved in UDCA degradation by *Sphingobium* sp. strain Chol11. Hsh3 eliminates the 7β-hydroxy group as water, leading to a double bond in the B-ring. This is similar to 7α-hydroxysteroid dehydratases in this and other strains, but Hsh3 is evolutionary different from these. Purified Hsh3 accepted steroids with and without side chain as substrates and had minor activity with 7α-hydroxy groups. The deletion mutant strain Chol11 Δ*hsh3* had impacted growth with UDCA and accumulated a novel compound. The compound was identified as 3’,5-dihydroxy-H-methyl-hexahydro-5-indene-1-one-propanoate, consisting of steroid rings C and D with a C_3_-side chain carrying the former 7β-hydroxy group, indicating that Hsh3 activity is important especially for the later stages of bile salt-degradation. Hsh3 homologs were found in other sphingomonads that were also able to degrade UDCA as well as in environmental metagenomes. Thus, Hsh3 adds to the bacterial enzyme repertoire for degrading a variety of differently hydroxylated bile-salts.

**Importance:** The bacterial degradation of different bile salts is not only important for the removal of these steroidal compounds from the environment but also harbors interesting enzymes for steroid biotechnology. The 7β-hydroxy bile salt UDCA naturally occurs in high concentration in the feces of black bears and has important pharmaceutical relevance for the treatment of different liver-related diseases, for which it is administered in high and increasing amounts. Additionally, it is present in the bile salt pool of humans in trace amounts. While ursodeoxycholate degradation is environmentally important, the enzyme Hsh3 modifies the hydroxy group that confers the medically relevant properties and thus might be interesting for microbiome analyses and biotechnological applications.

## Introduction

Bile salts are steroid compounds with different hydroxylation patterns that are produced by vertebrates. The dominant human bile salts all have a 3α-hydroxy group and can have additional 7α- and 12α-hydroxy groups. In contrast, ursodeoxycholate (UDCA, II in Fig 1) is a minor bile salt in humans (1), which has a 3α-hydroxy and a 7β-hydroxy group. It can be found as the dominant bile salt in Asian black bears (2, 3). Furthermore, UDCA is being used to treat different medical conditions (4, 5), especially liver diseases (6, 7) and gallstones (8, 9). For this, large amounts of UDCA (15 mg kg^-1^ body weight day^-1^) are administered, making up most of the bile salt pool during treatment (4, 10, 11) and requiring increasing production of UDCA. Currently, most of the UDCA is produced chemically and is not extracted from black bear bile anymore. New approaches of biotechnological UDCA production are subject to research (12, 13).

**Figure 1:**
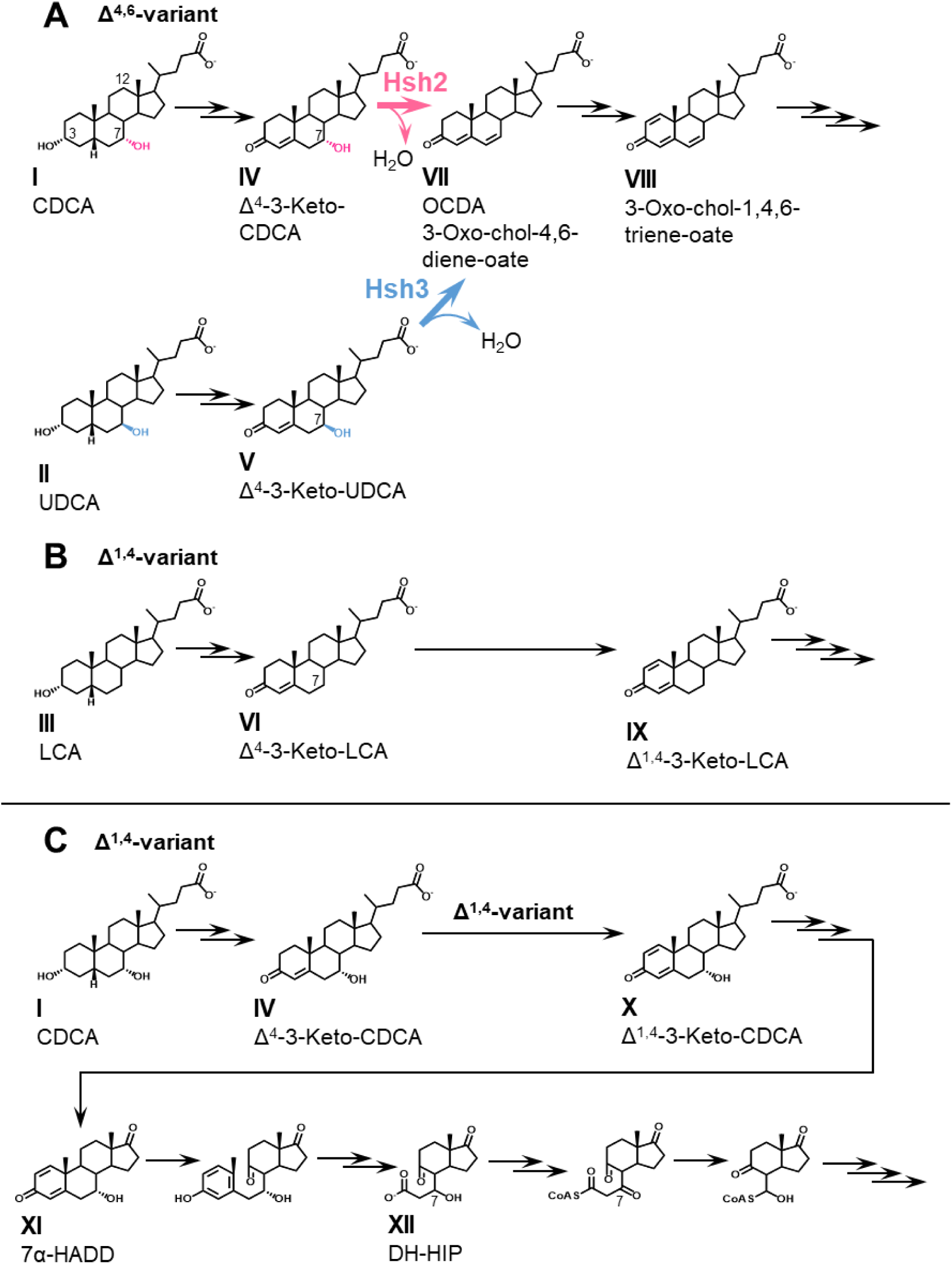
Degradation of bile salts in *Sphingobium* sp. strain Chol11 (A) and *Pseudomonas stutzeri* Chol1 (B+C). I, Chenodeoxycholate (CDCA); II, ursodeoxycholate (UDCA); III, lithocholate (LCA); IV, Δ^4^-3-keto-CDCA; V, Δ^4^-3-keto-UDCA; VI, Δ^4^-3-keto-LCA; VII, 3-oxo-chol-4,6-diene-oate (OCDA); VIII, 3-oxo-chol-1,4,6-triene-oate; IX, Δ^1,4^-3-keto-LCA, X, Δ^1,4^-3-keto-CDCA; XI, 7α-hydroxy-androsta-1,4-diene-3,17-dione; XII, 3′,7β-dihydroxy-3aα-*H*-4α(3′-propanoate)-7aβ-methylhexahydro-1,5-indanedione (DH-HIP).

While most bile salts are recycled in the so-called enterohepatic cycle, significant amounts are excreted (1, 14, 15). In the environment, bile salts can be degraded by bacteria via pathways similar to the degradation of other steroid compounds, namely the 9,10*-seco* pathway for aerobic degradation and the 2,3-*seco*-pathway for anaerobic degradation (15, 16) (Fig 1). One variant of bile salt degradation pathway (Δ^1,4^-variant), which can be found in *Pseudomonas stutzeri* Chol1 and *Comamonas testosteroni*, is well elucidated (17, 18) (Fig 1B). A second variant (Δ^4,6^-variant) can be found in sphingomonads such as *Sphingobium* sp. strain Chol11 and is only partly elucidated (15, 19, 20) (Fig 1A). Both pathway variants can be structured into four steps: bile salts are first oxidized to Δ^4^-3-keto bile salts (e.g., IV, V, VI in Fig 1; step 1: A-ring oxidation) (20–26). In step 2, side chain degradation yields C19-steroids (15, 27–30). In the Δ^1,4^-variant, androsta-1,4-diene-3,17-diones (ADDs, e.g., XI) are formed, which can be degraded aerobically via the 9,10-*seco* pathway initiated by monooxygenase-mediated cleavage of the B-ring (step 3: degradation of rings A and B) (17, 26, 31–34). A- and B-ring are degraded by *meta-*cleavage, hydrolytic and β-oxidation reactions (35–37), leading to derivatives of H-methyl-hexahydro-indanone-3-propenoate (HIP, e.g., XII). HIPs are central intermediates in both aerobic and anaerobic steroid degradation and are further degraded by β-oxidation and hydrolysis reactions (step 4: HIP degradation) (15, 27, 38, 39). Additional reactions channel the differently hydroxylated bile salts into common pathways. The 12α-hydroxy group is isomerized before ring cleavage reactions, and is then removed from HIP compounds (XII) by dehydration and reduction reactions (27, 28, 40). In contrast to this, the fate of the 7-hydroxy group differs depending on its conformation and the pathway. An hydroxy group at position of the former C7 is needed for the β-oxidation-like degradation of the former B-ring of HIP compounds (27). Therefore, a hydroxy group is introduced at this position in HIP if not present in the substrate bile salts by introduction of a double bond and hydration in *P. stutzeri* Chol1 and *C. testosteroni* (27, 41). While degradation of 7α-hydroxylated bile salts is well elucidated and found in many microorganisms, only few organisms have been described to degrade UDCA via this pathway, indicating an inhibiting effect of the 7β-hydroxy group (20, 42).

In the Δ^4,6^-variant, 7-hydroxy bile salt degradation proceeds differently (20): the 7-hydroxy group is eliminated as water leading to the formation of a double bond in the B-ring in Δ^4,6^-keto compounds prior to A-ring oxidation to Δ^1,4,6^-3-keto compounds (e.g., VII). The latter are the substrates for further degradation, including side chain removal via a yet largely unknown reaction sequence and most probably 9,10-*seco* cleavage (31). Genomic and proteomic studies strongly indicate that the degradation of the steroid nucleus is similar to the canonical degradation and also proceeds via HIP compounds (XII) (43). For 7α-hydroxy bile salts the dehydration reaction is catalyzed by 7α-hydroxy steroid dehydratase Hsh2 (20). A similar reaction catalyzed by a homologous enzyme, BaiE, can be found during the transformation of so-called primary 7α-hydroxy bile salts (e.g., chenodeoxycholate, CDCA, I) to secondary 7-deoxy bile salts by *Clostridium* members of the human gut microbiota (e.g., lithocholate, LCA, III) (44, 45). *Sphingobium* sp. strain Chol11 also degrades UDCA via the Δ^4,6^-pathway variant. However, Hsh2 does not catalyze 7β-hydroxy bile salt dehydration, and the deletion mutant still degrades UDCA via Δ^4,6^-3-keto compounds (VII), pointing at a second enzyme for 7β-dehydration (20). The 7β dehydratase reaction is also found during the transformation of UDCA to LCA by *Clostridia* in the gut during UDCA treatment. This reaction was hypothesized to be catalyzed by BaiI (14), but to the best of our knowledge, its function was not verified so far.

Therefore, our goal was to identify the 7β-hydroxy steroid dehydratase from the UDCA degradation pathway in *Sphingobium* sp. strain Chol11. As no other 7β-hydroxy steroid dehydratases were reported before, we attempted biochemical purification of this enzyme.

## Results

### The 7β-hydroxysteroid dehydratase in *Sphingobium* sp. strain Chol11 is encoded by a second gene and different from 7α-hydroxysteroid dehydratase Hsh2

As previously described, bile salt degradation intermediates with a Δ^4,6^-3-keto structure transiently accumulate in cultures of *Sphingobium* sp. strain Chol11 grown not only with the 7α-hydroxy bile salts cholate and CDCA, but also the 7β-hydroxy bile salt UDCA (20). While no Δ^4,6^-3-keto intermediates were formed from cholate and CDCA by strain Chol11 Δ*hsh2,* this strain still produced Δ^4,6^-3-keto intermediates from UDCA. Accordingly, a dehydratase activity towards the 7β-hydroxy substrate Δ^4^-3-keto-UDCA leading to formation of 3-oxo-chol-4,6-diene-oate (OCDA, VII in Fig 1) could be observed in cell extracts of *Sphingobium* sp. strain Chol11 as well as in cell extracts of strain Chol11 Δ*hsh2* (Fig 2), further confirming that this strain contains a second hydroxysteroid dehydratase that is able to transform 7β-hydroxy substrates. Surprisingly, the highest specific 7β-hydroxysteroid dehydratase activity was observed in the cell extracts of *Sphingobium* sp. strain Chol11 Δ*hsh2* grown with cholate, not UDCA. However, bioinformatical prediction of BaiI homologs did not lead to any active 7β-hydroxysteroid dehydratase (Text S1, Table S1), indicating the existence of a different type of hydroxysteroid dehydratase.

**Figure 2:**
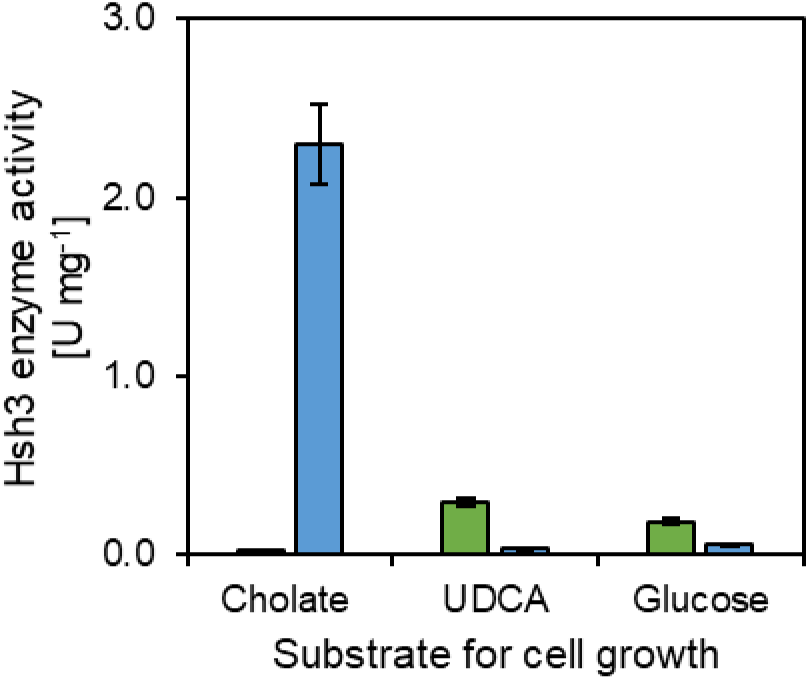
7β-Hydroxysteroid dehydratase activities in *Sphingobium* sp. strains Chol11 wt (green) and Δ*hsh2* (blue) cell extracts. Enzyme assays with extracts of cells grown with cholate, UDCA, or glucose as indicated, and Δ^4^-3-keto-UDCA (V in Fig 1) as substrate. All error bars indicate standard deviation of mean values (n=3).

### The 7β-hydroxysteroid dehydratase from *Sphingobium* sp. strain Chol11 is encoded by *nov2c681*

The same reverse-genetics approach of protein purification and identification by peptide mass fingerprinting as previously employed for the identification of Hsh2 (20) was used for identification of the 7β-hydroxysteroid dehydratase. As the highest 7β-hydroxysteroid dehydratase activity was found in the cell extracts of *Sphingobium* sp. strain Chol11 Δ*hsh2*, further protein purification was performed using cell extracts from this strain. An enzyme assay using Δ^4^-3-keto-UDCA (V in Fig 1, absorption maximum 245 nm) as substrate was used for all purification steps as activity could easily be monitored spectrophotometrically by the formation of the product OCDA (VII, absorption maximum at 290 nm). Final purification after ammonium sulfate precipitation and anion exchange chromatography was achieved by native PAGE (Fig S1), from which bands were excised and tested for 7β-hydroxysteroid dehydratase activity. The active band was subjected to peptide mass fingerprinting and identified as Nov2c681 (RefSeq ID WP_097093216.1).

In contrast to 7β-hydroxysteroid dehydratases Hsh2 and BaiE, Nov2c681 does not belong to the NTF2-like superfamily but has a dimeric α-β barrel EthD domain. Accordingly, it only has 16% and 12% identity to Hsh2 and BaiE from *C. scindens* VPI12708. It also has a low similarity (12% identity) to the putative 7β-hydroxy bile salt dehydratase BaiI from *C. scindens* ATCC35704. An AlphaFold3 prediction of the structure of Hsh3 indicated the presence of eight α-helices, which are organized in a nearly mirror-symmetrical manner around a barrel-like core of nine β-sheets (Fig 3). Overall, one major cavity was predicted (Fig 3A), which might represent an active site.

**Figure 3:**
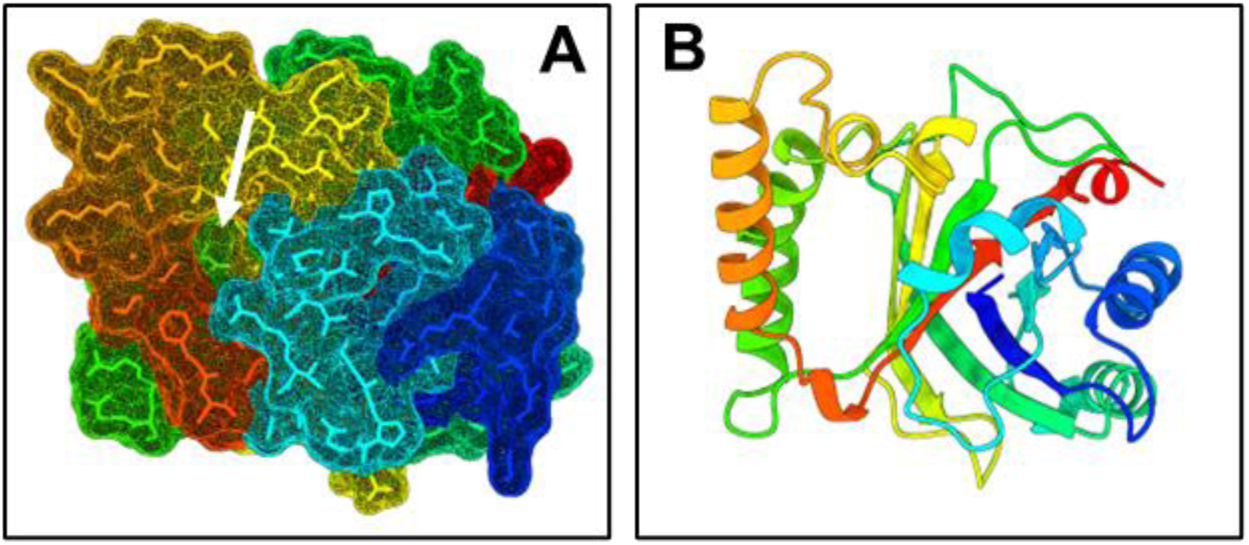
AlphaFold 3 prediction of the structure of Hsh3 from *Sphingobium* sp. strain Chol11. **(A)** All-atom representation with molecular surface. The white arrow indicates a cavity. **(B)** Ribbon diagram from the same angle as (A).

In previously described proteome analyses of *Sphingobium* sp. strain Chol11 (43), Nov2c681 was found in the proteomes of cells grown with all tested substrates cholate, deoxycholate, 7α,12β-dihydroxy ADD (7α,12β-DHADD), and glucose with higher amounts in the steroid-grown cells compared to the glucose control.

### Nov2c681/Hsh3 catalyzes the 7β-dehydration of ursodeoxycholate-derived compounds

For testing the activity of Nov2c681, *nov2c681* was heterologously expressed in *P. stutzeri* Chol1, resulting in 7β-hydroxysteroid dehydratase activity (Fig S2). *P. stutzeri* Chol1 pBBR1MCS-5::*nov2c681* transformed UDCA (II in Fig 1) to 3-hydroxy-9,10-*seco*-androsta-1,3,5(10),6-tetraene-9,17-dione (HSATD, see Fig S2) with a Δ^6^ double bond instead of 3,7β-dihydroxy-9,10-*seco*-androsta-1,3,5(10)-triene-9,17-dione (3,7β-DHSATD) that was produced by the wildtype strain (20). The production of HSATD also resulted in the production of a purple pigment that precipitated and could not be identified further, similar to the formation of a purple pigment from 3,12β-dihydroxy-9,10-*seco*-androsta-1,3,5(10),6-tetraene-9,17-dione (3,12β-DHSATD) at basic pH as reported before (46). Additionally, the enzyme was tested in *P. stutzeri* Chol1 Δ*kstD1* Δ*scdA1,* which transforms bile salts to their corresponding Δ^4^-3-keto derivatives. *P. stutzeri* Chol1 Δ*kstD1* Δ*scdA1* pBBR1MCS-5::*nov2c681* produced OCDA (VII) from UDCA, while CDCA (I) was transformed to Δ^4^-3-keto-Chenodeoxycholate (Fig S2). Thus, Nov2c681 has specific 7β-hydroxysteroid dehydratase activity, and it was renamed Hsh3 for **H**ydroxy**s**teroid de**h**ydratase 3.

### Hsh3 is active with different 7β-hydroxy steroids as well as minor activity with 7α-hydroxy bile salts

For characterizing Hsh3, especially in comparison with Hsh2, which has a very similar function, both enzymes were purified using a His-tag and affinity chromatography as well as gel filtration (Fig S3).

In contrast to most previous assays, kinetic parameters of the two enzymes were not determined in enzyme assays with Δ^4^-3-keto bile salts. Instead, the side chain-less compounds 7α- or 7β-hydroxy ADD (e.g., 7α-HADD, XI in Fig 1) for Hsh2 or Hsh3, respectively, were used, which were also transformed by these enzymes and allowed for more precise determination of transformation rates due to their absorption maximum at 300 nm (Fig S4, Table 3). The v_max_ of Hsh3 was about twice as high as the v_max_ of Hsh2, and Hsh3 had an about five-fold higher K_M_ compared to Hsh2, and a more than eight-fold higher k_cat_.

**Table 1:**
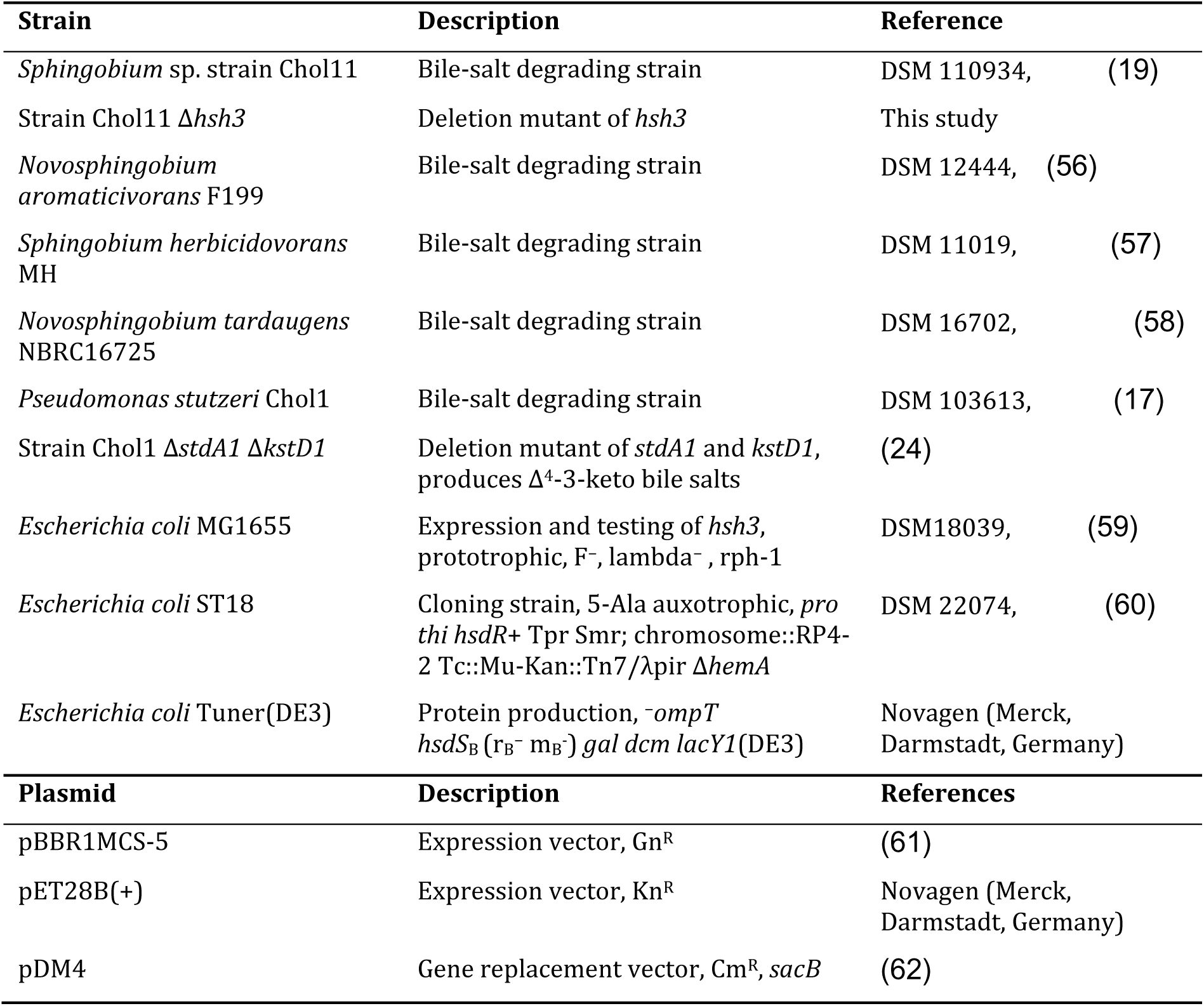
Strains and plasmids used in this study. Gn^R^, Kn^R^, CM^R^: Resistance against gentamicin, kanamycin, and chloramphenicol, respectively.

**Table 2:**
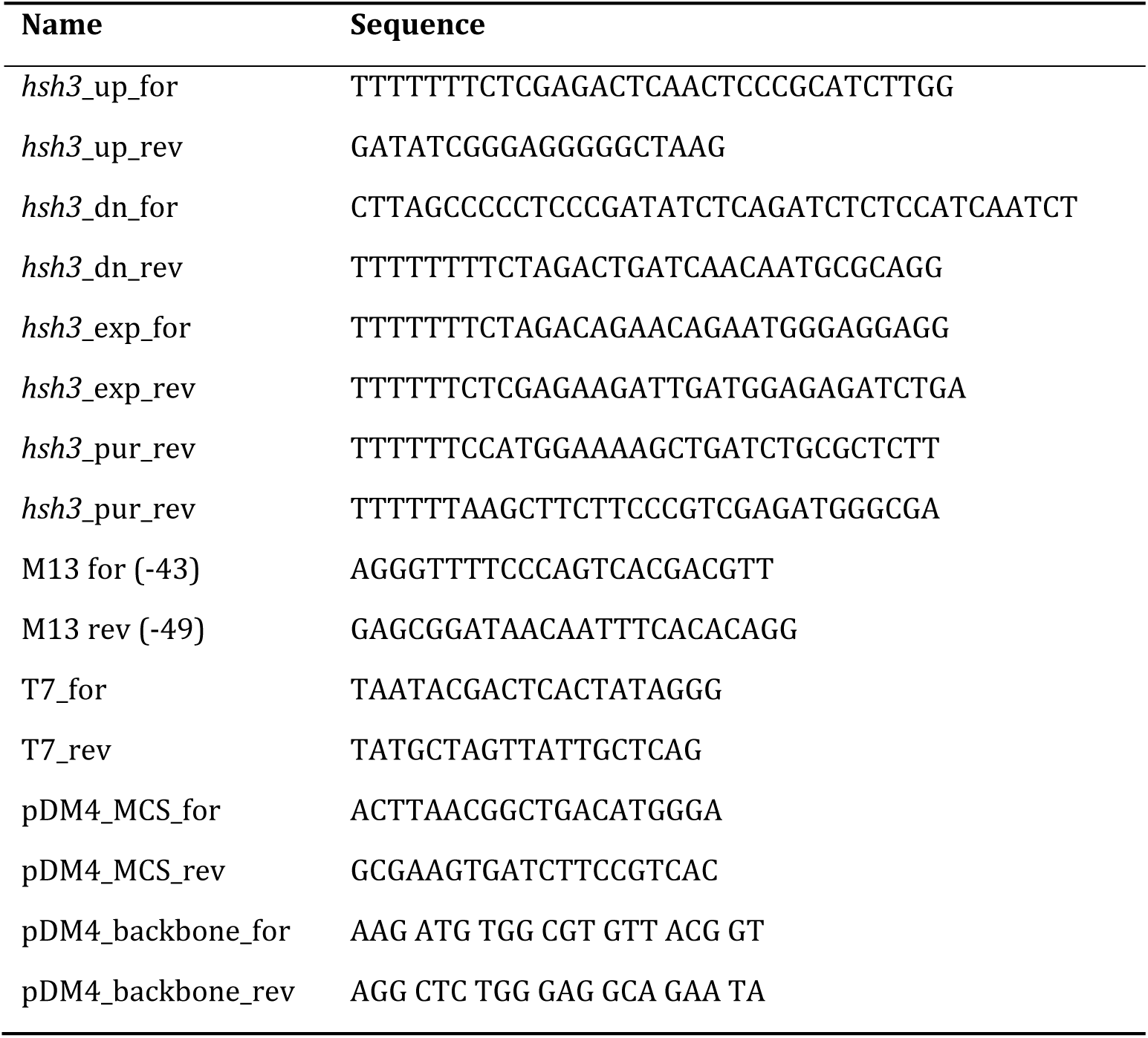
Primers used in this study.

**Table 3:**
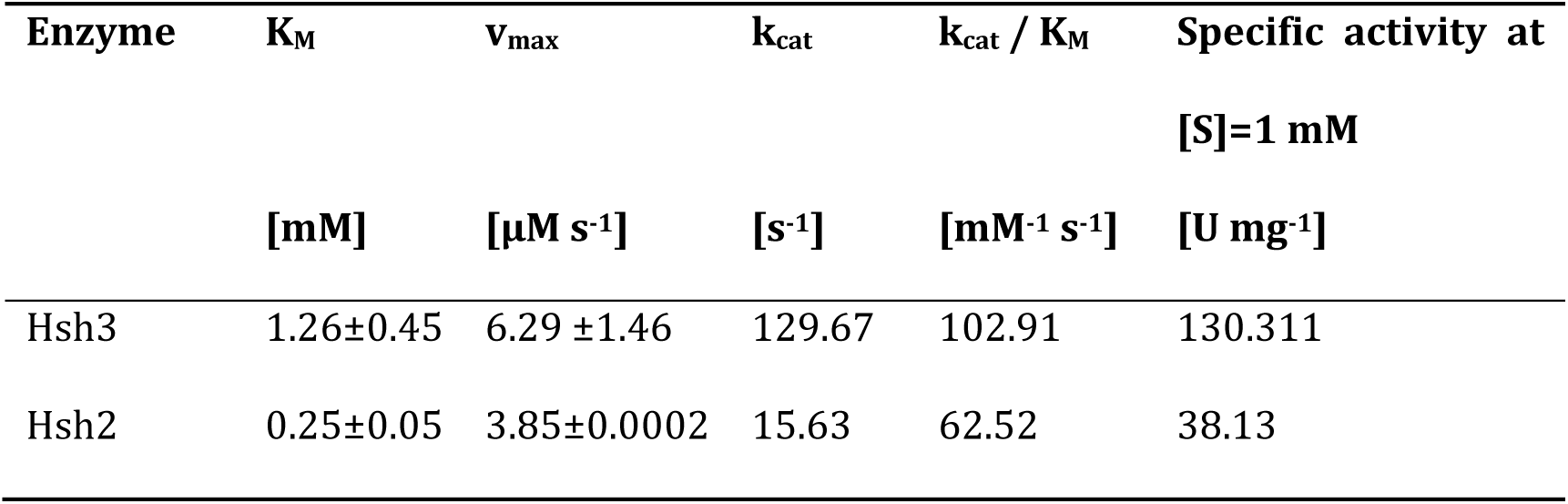
Kinetic parameters of purified Hsh2 and Hsh3 from *Sphingobium* sp. strain Chol11 as determined with 7α-HADD and 7β-HADD, respectively.

For further characterization, the ability to transform substrates with the respective other 7-hydroxy group was tested. When only 5 µg ml ^-1^ enzyme was added to the assay, only the respective substrates were transformed (Fig 4**Error! Reference source not found.**A+C). However, addition of much higher enzyme concentrations of 200 µg ml^-1^ led to the transformation of most of the added Δ^4^-3-keto-CDCA (IV) also by Hsh3, and limited transformation of Δ^4^-3-keto-UDCA (V) by Hsh2 (Fig 4B+D).

**Figure 4:**
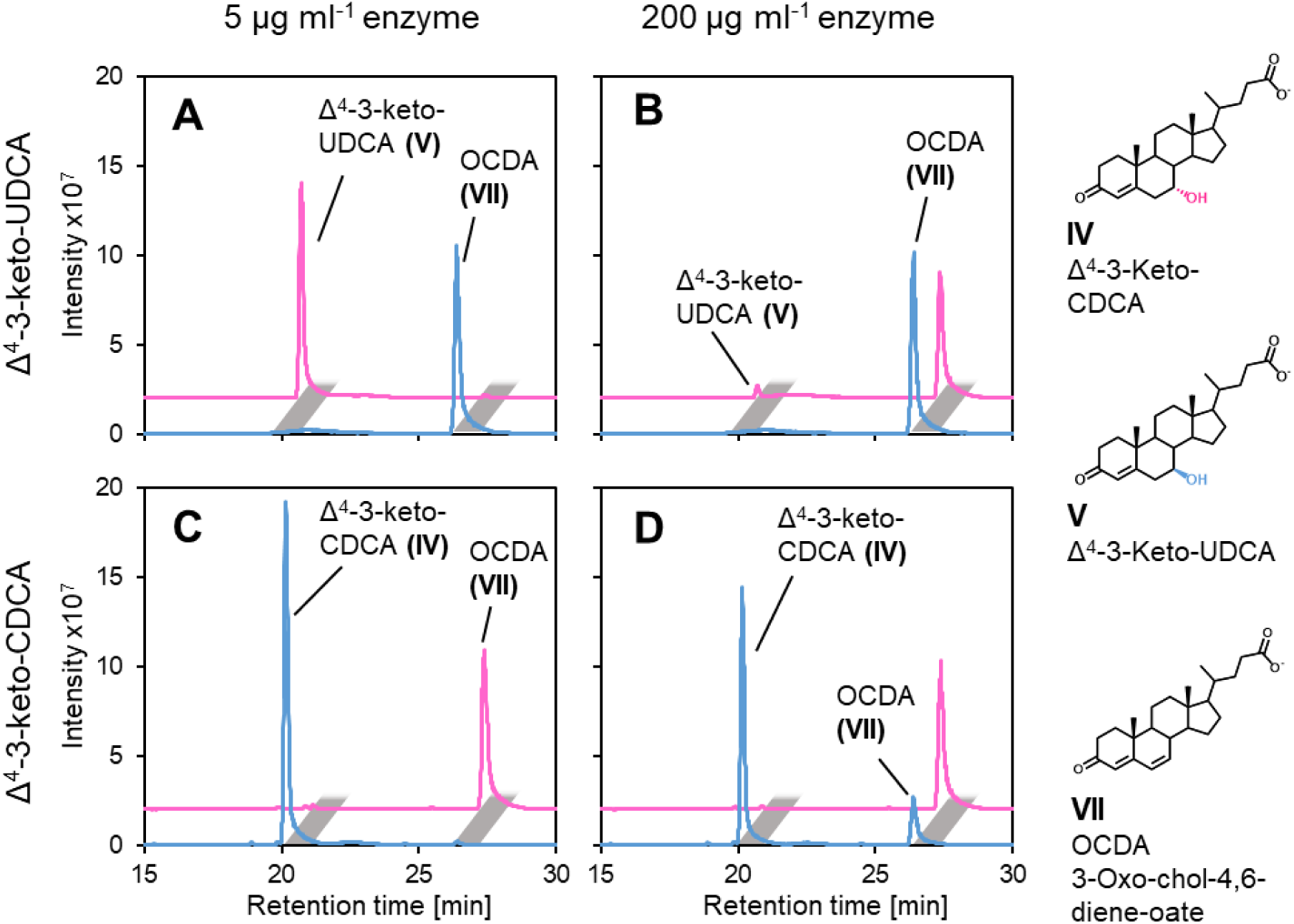
Transformation of Δ^4^-3-keto-UDCA (A+B) and Δ^4^-3-keto-CDCA (C+D) by purified Hsh3 (blue) and Hsh2 (red) from *Sphingobium* sp. strain Chol11 added in different concentrations (A+C, 5 µg ml^-1^; B+D, 200 µg ml^-1^). Incubated for 30 min at 30 °C. MS base peak chromatograms in negative mode.

### Hsh3 is involved in the degradation of 7β-hydroxy bile salt ursodeoxycholate in *Sphingobium* sp. strain Chol11 but not essential

For confirming the function of Hsh3 in UDCA degradation of *Sphingobium* sp. strain Chol11, the deletion mutant *Sphingobium* sp. strain Chol11 Δ*hsh3* was constructed. *Sphingobium* sp. strain Chol11 Δ*hsh3* displayed delayed growth and lower yields during growth with UDCA compared to growth with other bile salts. The lag phase increased to up to 30 h with UDCA, while no detectable lag phase was found with other bile salts, and the final OD_600_ that was about 15% to 30% lower than with other bile salts (Fig 5A). Plasmid-based expression of *hsh3* in the deletion mutant reversed the phenotype (Fig 5B). Expression of *hsh2* did not lead to wt-like growth of *Sphingobium* sp. strain Chol11 Δ*hsh3* (not shown).

**Figure 5:**
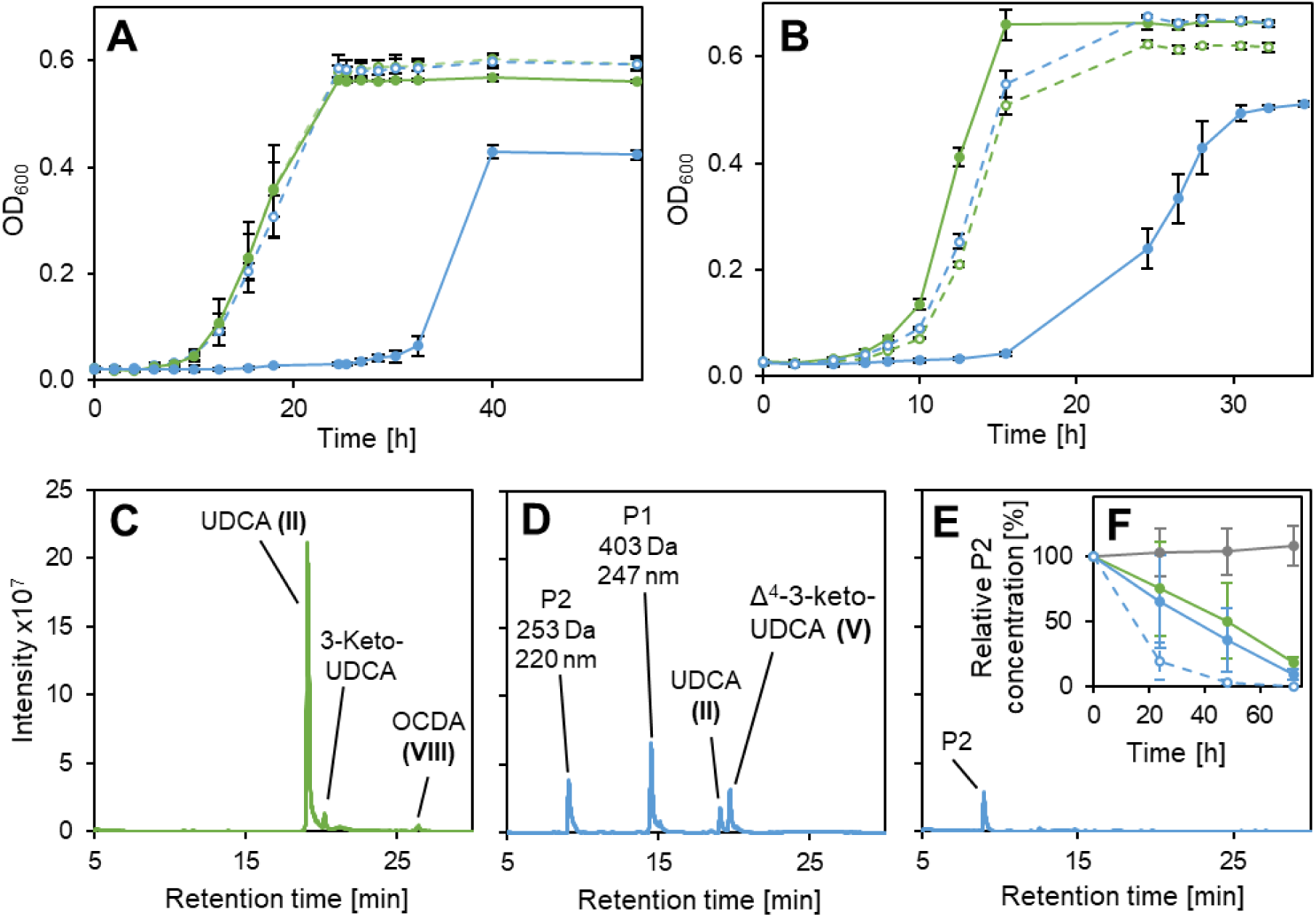
Growth of *Sphingobium* sp. strains Chol11 wt and Chol11 Δ*hsh3* with UDCA. **(A)** Growth of strain Chol11 Δ*hsh3* (blue) and wildtype (green) with either UDCA (solid lines) or CDCA (dashed lines). **(B)** Growth of strain Chol11 Δ*hsh3* (blue) and wt (green) with pBBR1MCS-5 (solid lines) or with pBBR1MCS-5::*hsh3* (dashed lines) with UDCA. **(C)** Transient accumulation of Δ^4,6^-3-keto compound OCDA during degradation of UDCA by strain Chol11 wildtype. **(D)** Accumulation of Δ^4^-3-keto UDCA as well as two new compounds (P1 and P2) by strain Chol11 Δ*hsh3* during degradation of UDCA. **(E)** Accumulation of product P2 in cultures of strain Chol11 Δ*hsh3* in stationary phase after 26 h. Supernatants from exponential phase (C,D) or stationary phase (E) cultures. MS base peak chromatograms in negative mode. **(F)** Degradation of P2 by strain Chol11 wt (green) or Δ*hsh3* (blue). Grey, sterile control; solid lines, OD_600_=1; dashed lines, OD_600_=2. Cell suspensions were incubated at 30°C. All error bars indicate standard deviation of mean values (n=3).

### *Sphingobium* sp. strain Chol11 Δ*hsh3* produces a HIP compound that can only slowly be degraded

During growth with UDCA, *Sphingobium* sp. strain Chol11 Δ*hsh3* accumulated several additional compounds compared to the wildtype, but no Δ^4,6^-3-keto (e.g., VII in Fig 1) compounds as indicated by the respective UV spectra (Fig 5D). During the exponential growth phase, some remaining UDCA (II) as well as Δ^4^-3-keto-UDCA (V) were found. Additionally, one so-far unknown compound P1 with a mass of 403 Da and an absorption maximum at 247 nm was observed. The absorption spectrum indicates a Δ^4^- or Δ^1,4^-3-keto structure of the steroid skeleton. Interestingly, the 16 Da higher mass compared to Δ^4^-3-keto-UDCA indicates a hydroxylation at a so-far unknown position. Another compound, P2, was present in these cultures and remained in the supernatant even after several days of incubation (Fig 5E). This compound had a molecular mass of 253 Da and an absorption maximum at 220 nm (Fig 6A+B). The culture of the deletion mutant was slightly more yellow compared to the wildtype cultures, which was not due to cell color but the supernatant (Fig S5A). For determining if P2 could be a natural degradation intermediate of *Sphingobium* sp. strain Chol11 or a modified side product, degradation of the compound by strain Chol11 was tested. Interestingly, both, strain Chol11 wt and strain Chol11 Δ*hsh3,* were able to degrade over 80% of the added amount within several days (Fig 5E). Biotic degradation was confirmed by a sterile control, in which no degradation was observed, and a control containing twice as many cells, in which degradation was about twice as fast. The mass of compound P2 indicated a structure similar to HIP compounds (e.g., XII). For analyzing P2 in more detail, it was purified by extraction, which resulted in a yellow solution (Fig S5B), and HPLC purification. P2 was acid-but not base-stable (Fig S5C-E**Error! Reference source not found.**). The structure of P2 was analyzed by NMR measurements (Fig S6). Although P2 was purified by HPLC, two distinct structures P2A and P2B with different molecular masses were identified by NMR (Fig 6C). The mixture contained 75% compound P2A and 25% compound P2B. As predicted, P2A is a HIP compound, 3’,5-dihydroxy-H-methyl-hexahydro-5-indene-1-one-propanoate (3,5-DH-HIEP), which still has the former 7β-hydroxy group. In addition, it has a double bond in the former C-ring and the former 9-keto group is reduced to a hydroxy group, which probably is a result of keto-enol tautomerism. The second product P2B has a lactone ring and might be formed from P2A by cyclization and water loss at former C7. Additionally, the compound has a different double bond in the former C-ring, which might also be a product of the keto-enol tautomerism.

**Figure 6:**
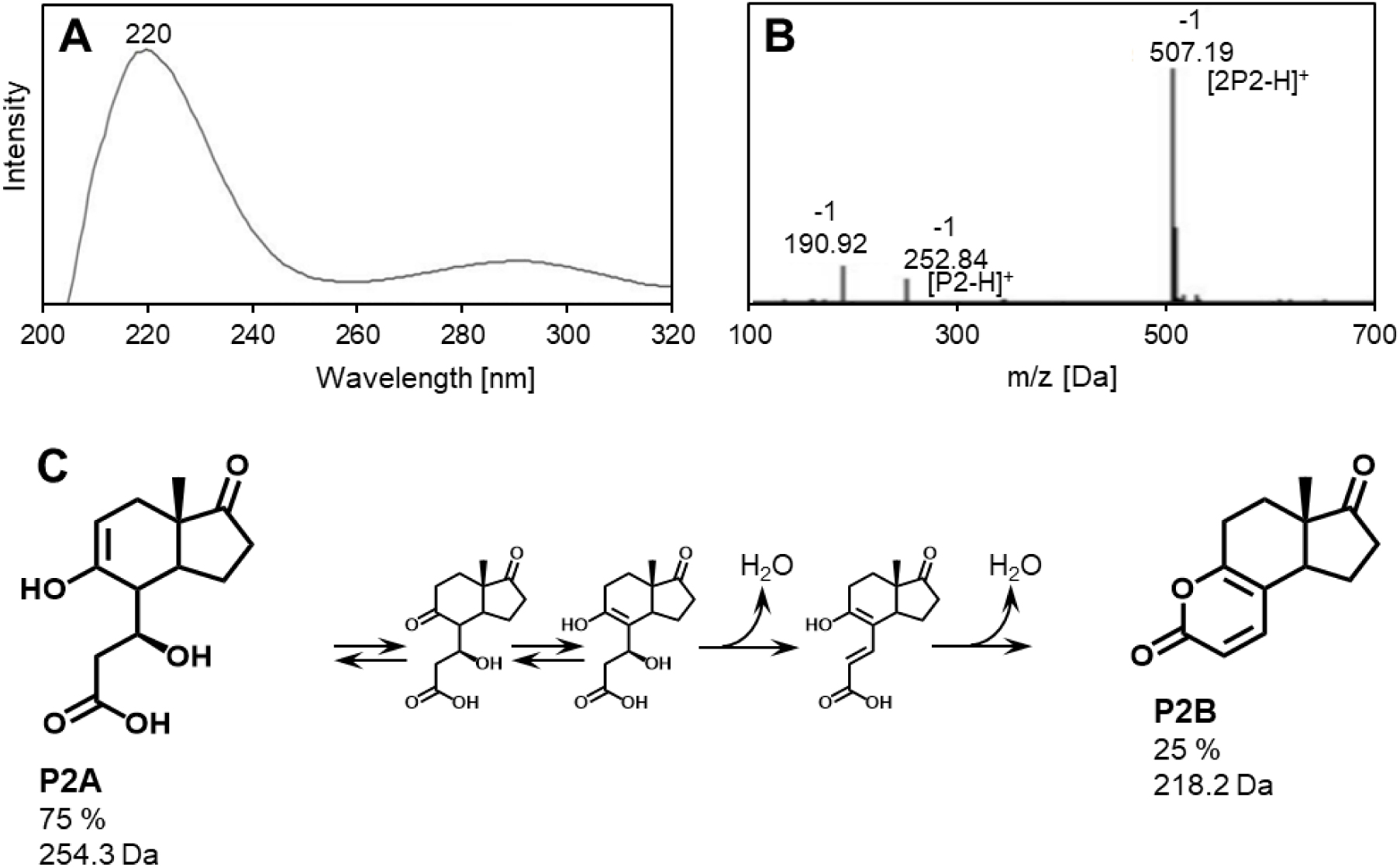
Characteristics and structure of compound P2 produced by *Sphingobium* sp. strain Chol11 Δ*hsh3* from UDCA. **(A)** UV spectrum of P2. **(B)** Mass spectrum of P2. **(C)** Structures of the compounds P2A and P2B found in the extracted P2 solution including a potential mechanism for the abiotic formation of P2B from P2A.

### Hsh3 homologs and activity can be found in several bile salt-degrading *Sphingomonadaceae* strains, further expanding the metabolic repertoire for bile salt in these strains

Recently, bile salt degradation via the Δ^4,6^-pathway was predicted for many strains of the *Sphingomonadaceae* and confirmed for three of these strains, *Sphingobium herbicidovorans* MH, *Caenibius tardaugens* NBRC16725, and *Novosphingobium aromaticivorans* F199 (43). A reciprocal BLASTp analysis showed that all three strains contain one single homolog of Hsh3 with very high identities of at least 62 % to Hsh3 (Table 4). Growth experiments confirmed growth of *N. aromaticivorans* F199 and *C. tardaugens* NBRC16725 with UDCA and a transient accumulation of Δ^4,6^-intermediates during growth (Fig 7).

**Figure 7:**
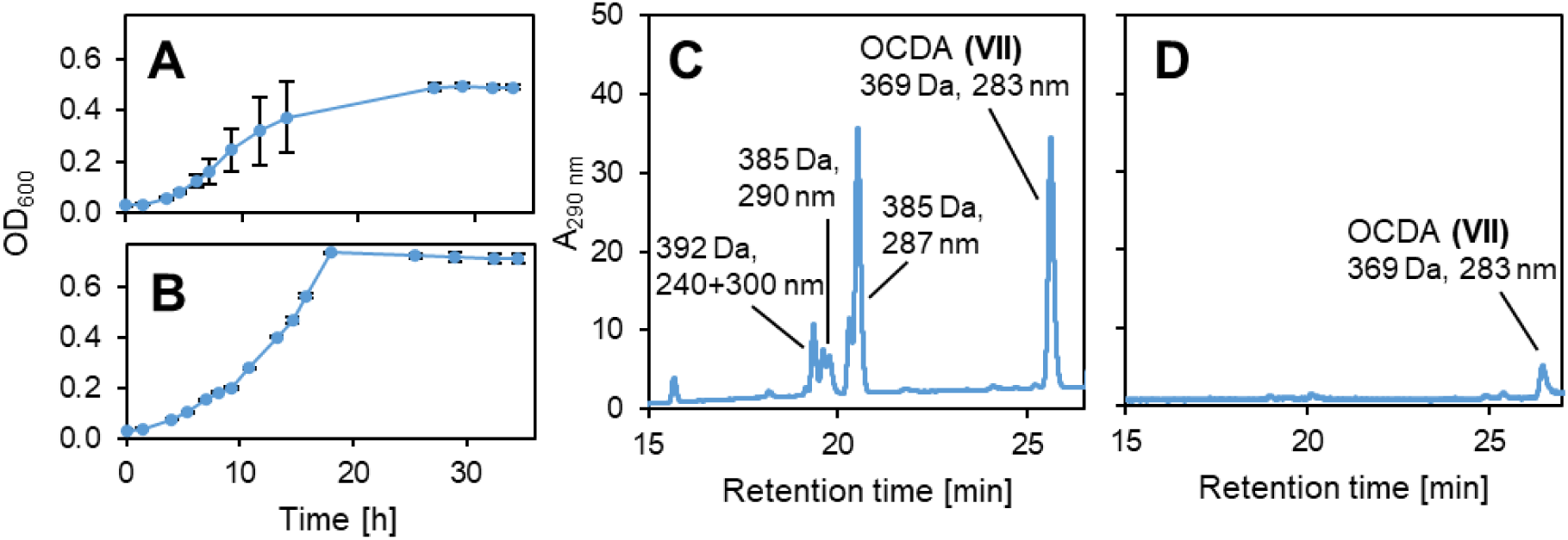
Growth of other sphingomonads with UDCA and accumulation of Δ^4,6^-3-keto compounds. (A+C) *Novosphingobium aromaticivorans* F199. **(B+D)** *Caenibius tardaugens* NBRC16725. (A+B) Growth with 1 mM UDCA. (C+D) MS base peak chromatograms in negative mode. All error bars indicate standard deviation of mean values (n=3).

**Table 4:** Hsh3 homologs in other bile salt-degrading strains of the *Sphingomonadaceae* as determined by BLASTp analysis. RefSeq accession numbers are given as identifiers.

| Strain | Hsh3 homolog | Identity | E-value |
| --- | --- | --- | --- |
| *S. herbicidovorans* MH | WP_234810817.1 | 84 % | 1e-117 |
| *N. aromaticivorans* F199 | WP_011906908.1 | 62 % | 6e-101 |
| *C. tardaugens* NBRC16725 | WP_021690432.1 | 64 % | 2e-112 |

### Ursodeoxycholate-degrading bacteria are present in a wide variety of environments with bile salt input

Using an HMM search on the MGnify website, homologs for Hsh3 as well as Hsh2 could be found in metagenomes from many environments, especially from environments classified as environmental - marine and engineered – wastewater (Fig S8). The highest fraction of assemblies with Hsh3 and Hsh2 homologs was from aquaculture environments. Strikingly, only very few homologs were found in metagenomes from host-associated environments, and none in metagenomes from human digestive systems.

For finding further UDCA degraders, and finding potential other pathways for UDCA degradation, enrichment cultures with UDCA were inoculated from different sites with bile salt input such as duck ponds and soil close to agricultural fields. This way, many UDCA-degrading strains could be isolated, pointing at a wide distribution of this capacity in the environment. According to their 16S rRNA sequences, some strains belonged to the *Sphingomonadaceae,* but most strains belong to the groups *Pseudomonas* or *Comamonas* (Fig S9). As expected, the *Sphingomonadaceae* strains transiently accumulated Δ^4,6^-intermediates. However, some *Pseudomonas* and *Comamonas* strains were also able to completely degrade UDCA without transient accumulation of Δ^4,6^ compounds and only produced Δ^1,4^ compounds indicating the existence of an additional pathway.

## Discussion

With Hsh3, we found an additional enzyme modifying the diverse hydroxy groups of bile salts and describe the first 7β-hydroxysteroid dehydratase. The different hydroxylation patterns of bile salts among different animals lead to an astonishing diversity of these compounds (47). At the same time, many bacteria interacting with these compounds have evolved enzymes to modify bile salts at their hydroxy groups: On the one hand, intestinal bacteria oxidize and eliminate these hydroxy groups as well as change their conformation (48). On the other hand, environmental bacteria degrade bile salts as energy and carbon sources and modify the hydroxy groups to allow degradation via unified pathways (15). Our data shows that 7β-hydroxysteroid dehydratase Hsh3 catalyzes the dehydration of the 7β-hydroxy group of UDCA, thus allowing *Sphingobium* sp. strain Chol11 to channel this bile salt into the same pathway as CDCA and CA, which are dehydrated by the 7α-counterpart Hsh2 (20). Its homologs in various sphingomonads add to the wide bile salt substrate spectrum of these bacteria (43). UDCA-degrading strains transiently accumulating Δ^4,6^-steroid intermediates could also be isolated from lakes and agricultural soils in Germany. Additionally, we found homologs of Hsh3 in many wastewater treatment plant and aquatic microbiomes, especially aquaculture microbiomes, which are probably also contaminated with bile salts. Together, this indicates a wide distribution of the Δ^4,6^-degradation pathway and points at the widespread presence of 7β-hydroxysteroid compounds in these environments.

Despite its role in UDCA degradation in *Sphingobium* sp. strain Chol11, Hsh3 does not seem to be indispensable for UDCA degradation, with deletion of *hsh3* only leading to delayed growth and decreased yields. As expected, no Δ^4,6^-intermediates were observed in the supernatants of UDCA cultures. This is similar to *Sphingobium* sp. strain Chol11 Δ*hsh2*, which is also able to grow with 7α-hydroxy bile salts without forming Δ^4,6^-intermediates (20). However, *Sphingobium* sp. strain Chol11 Δ*hsh3* accumulates two new intermediates when grown with UDCA, which highlights bottlenecks in degradation by this strain and might explain the lower final OD_600_. Intermediate P1, which is only transiently accumulated, has a Δ^1,4^- or Δ^4^-3-keto structure, like the other intermediates found in this culture. However, the mass of P1 indicates one additional hydroxylation compared to UDCA, the position of which is unclear. Other hydroxylated compounds have been found in the supernatants of *Sphingobium* sp. strain Chol11 Δ*sclA* and Δ*scd4A*, which carry deletions of side chain degradation genes (31). On the one hand, the hydroxylation could be at C9, a reaction necessary for later B-ring cleavage, which is catalyzed by KshA homologs Nov2c407, Nov2c430, and Nov2c440 in *Sphingobium* sp. strain Chol11. On the other hand, it might be in another position and important for side chain degradation. This hydroxylation is probably catalyzed by the monooxygenase Nov2c228 in *Sphingobium* sp. strain Chol11 (31). As P1 and other intermediates with side chain are subsequently degraded by *Sphingobium* sp. strain Chol11 Δ*hsh3*, the β-hydroxy group seems to slow down degradation during or after side chain degradation. The other new intermediate, P2A, remained in the culture supernatant after the culture reached stationary phase, but could be degraded by both the wildtype strain and *Sphingobium* sp. strain Chol11 Δ*hsh3*. By NMR spectroscopy P2A was shown to be a HIP-compound consisting of only the C- and D-ring with a C_5_-side chain from the former B-ring. As expected, the former 7β-hydroxy group is still found in P2A. An unusual enol structure in P2A is most probably stabilized by an intramolecular hydrogen bond between the enol hydroxy group and the carboxy group, as indicated by NMR spectroscopy (Fig S10). A second structure P2B found together with P2A after purification could be derived from P2A abiotically by dehydration, which has also been observed for steroid compounds under acidic conditions as used for extraction (49) and lactone formation, which is has also been observed before for these HIP compounds (50, 51). For *C. testosteroni*, it was suggested that degradation of HIP compounds from steroids involves a β-hydroxy group in the former C7 position. During the degradation of 7α-hydroxy steroids such as cholate and CDCA, this would be achieved by consecutive dehydration (similar to Hsh2 reaction, but later in degradation) and hydration in β-conformation (52). Therefore, it might be speculated that the Hsh2 reaction in sphingomonads is a preparation for this step, especially as this reaction can be observed as a minor - potentially side - reaction in *C. testosteroni* as well (40). However, degradation of the β-hydroxy HIP (P2A) was strongly delayed in *Sphingobium* sp. strain Chol11 Δ*hsh3*. Additionally, *Sphingobium* sp. strain Chol11 Δ*hsh2* did not accumulate similar HIP compounds as persistent intermediates. This indicates that, at least in *Sphingobium* sp. strain Chol11, a 7α-hydroxy group is needed for β-oxidation of the HIP side chain. In either case, the hydroxy group is first eliminated by Hsh2 or Hsh3, and then the hydroxy group in a suitable conformation is added at a later stage. This could indicate either a strategy to funnel all bile salts into the same pathway, or point at enzymes that could be inhibited by these hydroxy groups.

Hsh3 and Hsh2 displayed minimal activity with the respective other substrate. However, this minimal activity was not enough to reverse the phenotype of *Sphingobium* sp. strain Chol11 Δ*hsh3*, even if *hsh2* was overexpressed from an expression vector. At the same time, the side activity of the two enzymes could explain the high Hsh3 activity in the *Sphingobium* sp. strain Chol11 Δ*hsh2* cell extract: *hsh3* might be overexpressed to compensate for the *hsh2* deletion.

Interestingly, Hsh3 is very different from Hsh2 despite catalyzing very similar reactions, indicating different evolutionary origins. Additionally, Hsh3 is structurally different from the proposed 7β-hydroxysteroid dehydratase BaiI from *C. scindens* and Hsh3 homologs are not present in *C. scindens.* This points at the existence of phylogenetically distant additional 7β-hydroxy dehydratases in *Clostridia,* which is further emphasized by the lack of Hsh3 homologs in *Clostridium* genomes according to our BLAST search. Additionally, we did not find any Hsh3 homologs in human digestive system metagenomes in the MGnify database or in the metagenomes of the human microbiome project (53). However, we also did not find any Hsh2 homologs in these metagenomes, despite the high similarity of Hsh2 and BaiE, which indicates that our analysis might be too specific to find any more distant homologs in metagenomic datasets.

Besides its relevance for the degradation of UDCA, which is probably released into the environment in large amounts due to its pharmaceutical use, Hsh3 could be of interest for biotechnological applications for production of tailored bile salts. Hydroxysteroid dehydratases Hsh2 and Hsh1 are similar enzymes but catalyze the elimination and addition of hydroxy groups in steroidal compounds, respectively. Therefore, a reverse Hsh3 reaction producing UDCA-like compounds seems possible and interesting.

## Materials and methods

### Bacterial strains and cultivation

Bacterial strains were cultivated as described previously (31) and strains and plasmids used in this study are listed in Table 1. Strains of *E. coli* were grown in lysogeny broth medium (LB, (54)) if not described otherwise. Strains of *Sphingobium* sp. strain Chol11 and *Pseudomonas stutzeri* Chol1 were grown in the HEPES buffered medium B (MB, (55)) with 1 mM bile salts, 12 mM succinate (strain Chol1), or 15 mM glucose (strain Chol11) as described. *Novosphingobium aromaticivorans* F199 and *Novosphingobium tardaugens* NBRC16725 were cultivated in MB with 1 mM bile salt as carbon source. *E. coli* strains were cultivated at 37 °C on agar plates, and all other strains and *E. coli* strains in liquid cultures were cultivated at 30 °C.

For strains containing pBBR1MCS-5 20 µg ml^-1^ gentamicin was added. For strains containing pDM4 30 (*E. coli*) up to 90 (strain Chol11) µg ml^-1^ chloramphenicol was added, and for strains containing pET28B(+) 50 µg ml^-1^ kanamycin was added. For cultivation of *E. coli* ST18, 50 µg ml^-1^ 5-aminolevulinic acid was added. For agar plates, 1.5 % bacto agar (BD, Sparks, USA) were added to the respective media.

For pre-cultures, 5 ml medium in test tubes was inoculated from agar plates and incubated at 30 °C and 200 rpm overnight (ca. 14 h). For growth experiments, 5 ml cultures in test tubes were inoculated at an initial optical density at 600 nm (OD_600_) of about 0.02 using pre-cultures and incubated at the same conditions. Growth was tracked by measuring OD_600_. HPLC-MS samples were withdrawn at regular intervals and supernatants were either analyzed directly or stored at −20 °C.

### Biotransformation experiments

For transformation of steroid compounds with different bacteria expressing *hsh3* and the respective empty vector controls, cells from pre-cultures were washed and resuspended in MB medium at the given OD_600_. Steroids as well as the respective non-steroid carbon source were added to these cell suspensions before incubation at 30 °C. Samples for HPLC-MS measurements were withdrawn after addition of the steroid substrate as well as at further time points, stored at −20 °C, and centrifuged prior to HPLC-MS analysis of the supernatant.

### Enrichment, isolation and characterization of ursodeoxycholate-degrading strains

Enrichment cultures for the isolation of UDCA-degrading strains were prepared in MB as described (19) and inoculated from environmental samples. Sampling sites included duck ponds, lake Aasee, as well as soil and water from drainage of agricultural fields in the Münsterland area.

1 ml of water samples was combined with 4 ml MB with 1 mM UDCA and incubated in test tubes at 30 °C and 200 rpm. For soil samples, 1 g soil was suspended in 5 ml MB at room temperature and 200 rpm for 1 h. 1 ml of these suspensions was added to 4 ml MB with 1 mM UDCA and diluted 1:10 in the same medium before incubation at 30 °C and 200 rpm. Activity in these cultures was tracked by OD_600_ and HPLC-MS measurements of the supernatant. Turbid cultures were transferred to fresh medium for three times before transfer to agar MB agar plates with 1 mM UDCA. Single colonies with different morphology were transferred to new plates three times for isolation of single strains.

For identification of isolates, genomic DNA was extracted from the cells and the 16s rDNA was sequenced after amplification. Strains were then identified using BLASTn (63).

### Cloning techniques and construction of an unmarked deletion mutant

Cloning was performed according to standard procedures and as described (24). Primers for the cloning procedures are listed in Table.

For expression of proteins in different strains, the respective gene was amplified from genomic DNA using the exp_for and exp_rev primers. The gene and expression plasmid pBBR1MCS-5 were processed using the given restriction enzymes (Table), ligated, and transferred into *E. coli* MG1655 or ST18 by heat shock transformation. Plasmids were transferred into strains Chol1 or Chol11 by bi-parental conjugation from *E. coli* ST18. Correct assembly of vectors and transformation were confirmed by colony PCRs using primers M13 and sequencing.

For construction of deletion mutants, up- and downstream fragments of the respective gene were amplified using primer pairs up_for/rev and dn_for/rev, respectively. The fragments were fused using splicing by overlapping extension (SOE)-PCR (64) using the primer pair up_for/dn_rev and cloned into vector pDM4 using the given restriction enzymes. Correct vector assembly was confirmed using primer pair pDM4_MCS_for/rev after transformation into *E. coli* ST18. The resulting vector was transferred into strain Chol11 by conjugation and successful conjugation and insertion was confirmed by colony PCR using primer pair pDM4_backbone_for/rev. Unmarked gene deletion after forcing vector excision by cultivation on plates containing 10% sucrose was confirmed by colony PCR with primer pair up_for/dn_rev and sequencing.

### Enrichment, isolation and identification of Hsh3

Hsh3 was purified from cell extracts of UDCA-grown cells of *Sphingobium* sp. strain Chol11 as described for Hsh2 by (20).

For this, four 500 ml cultures of strain Chol11 Δ*hsh2* in MB with 1 mM cholate in Erlenmeyer flasks were inoculated from pre-cultures. Cells of each culture were harvested in the exponential growth phase by centrifugation at 4 °C and 4000 xg for 30 min, washed with 50 mM MOPS buffer (pH 7.8), and resuspended in 4 ml 50 mM MOPS buffer (pH 7.8). Cells were then lysed by ultrasonication (UP200S, Hielscher Ultrasonics, Teltow, Germany) with amplitude 60% and cycle 0.5 for 14 min on ice with 1 min breaks every 4 min. Cell debris was removed by centrifugation at 4 °C and 25000 xg for 30 min, and cell-free extracts from all cultures were combined for further isolation steps.

The first purification step was ammonium sulfate precipitation. After prior optimization, 40% (w/v) ammonium sulfate was added to the cell extracts in a first step. Precipitated proteins were removed by centrifugation at 0 °C and 25000 xg for 12 min. In a second step, the ammonium sulfate concentration in the remaining supernatant was increased to 60%, and precipitated proteins were again removed by centrifugation. The supernatant was discarded and the precipitate of the second step was used for further purification. For this, the precipitate was solved in 50 mM MOPS buffer (pH 7.8) and desalted by gel filtration using PD-10 desalting columns (GE Healthcare, Chicago, Il, USA) as recommended. For equilibration and elution, 3.5 ml 20 mM MOPS buffer (pH 7.0) was used. The resulting protein solution was further separated by anion exchange chromatography using a SOURCE 15Q column (6 ml, GE Healthcare, Chicago, Il, USA). After equilibration with 20 mM MOPS buffer (pH 7.0), the sample was loaded onto the column, and the column was washed with 20 volumes of the same buffer and inverted. For elution, a linear gradient towards a final concentration of 0.7 M NaCl in the same buffer over 20 column volumes was applied and 1 ml fractions were collected.

Fractions were tested for 7β-hydroxysteroid dehydratase activity and active fractions were subjected to native PAGE. Gels consisted of a gradient from 7% over 9% to 15% acrylamide (acrylamide/bisacrylamide 37.5:1) in Tris-HCl buffer (final concentration 325 mM, pH 8.8) with 0.2 % ammoniumperoxodisulfate and 0.02% N,N,N’,N’-tetramethylendiamine. 25 mM Tris with 192 mM glycine at pH 8.2 was used as running buffer and 100 mM Tris-HCl (pH 6.8) with 50% glycerol was used as loading buffer. Samples were loaded onto the gel twice and symmetrically, so that the gel could be cut in to two identical halves after separation. One half was stained using Coomassie blue to identify protein bands and used as an orientation to cut the corresponding areas from the second, unstained half of the gel.

Protein bands were then incubated in 800 µl 50 mM MOPS buffer (pH 7.8) at room temperature and 1200 rpm for 30 min. Supernatants were then used for finding bands with 7β-hydroxysteroid dehydratase activity. Protein bands from the stained gel corresponding to bands with activity were then analyzed by peptide mass fingerprinting.

### Purification of Hsh3 and Hsh2

For purification of Hsh3 using a his-tag, *hsh3* was cloned into the vector pET28(+)B using primer pair pur_for/rev for amplification and primer pair T7_for/rev for sequencing. pET28(+)B::*hsh3* was transferred into *E. coli* Tuner(BL3).

Two 250 ml cultures of each *E. coli* Tuner(BL3) pET28(+)B::*hsh3* (this study) and *E. coli* Tuner(BL3) pET28(+)B::*hsh2* (20) were inoculated in Erlenmeyer flasks from the respective pre-cultures at an initial OD_600_ of 0.02 and incubated at 37 °C and 120 rpm for 2 h. 0.2 mM IPTG were added and the cultures were further incubated at room temperature and 120 rpm overnight. Cells were harvested by centrifugation at 4 °C and 8000 xg for 8 min and kept on ice. Cells were washed and resuspended in 10 ml phosphate-buffered saline and lysed by ultrasonication (UP200S, Hielscher Ultrasonics, Teltow, Germany) with amplitude 60% and cycle 0.5 for 10 min on ice with 1 min breaks every 4 min. Cell debris was removed by centrifugation at 4 °C and 20000 xg for 60 min. Protein concentration in the cell-free extract was determined by BCA assay (Pierce, Thermo Fisher Scientific, Waltham, Ma, USA).

Hsh3 and Hsh2 were purified using HisPur Ni-NTA spin columns (0.2 ml, Thermo Fisher Scientific, Waltham, Ma, USA) according to the instructions. Column were loaded with cell-free extracts, adding a total protein amount corresponding to twice the binding capacity of the column in two steps. Protein containing fractions with concentrations >10 mg ml^-1^ were further purified by gel filtration for imidazole removal. Gel filtration was performed using HiTrap desalting columns (Cytiva, Marlborough, Ma, USA) according to the manual “operation with syringe” using 20 mM Tris buffer (pH 8) containing 150 mM NaCl. Fractions containing the protein were combined.

All purification steps were confirmed using SDS-PAGE.

### Enzyme assays

Enzyme assays were either conducted with purified Hsh3 and Hsh2 or with cell-free extracts. Cell-free extract were prepared as described (24). Strains were grown in 50 ml cultures in Erlenmeyer flasks and harvested in the exponential phase by centrifugation at 4 °C and 8000 xg for 10 min. The cells were washed and resuspended in 1.5 ml 50 mM MOPS buffer (pH 7.8), and lysed by ultrasonication (UP200S) with amplitude 60% and cycle 0.5 for 8 min on ice with 2 min after the first 4 min. Cell debris was removed by centrifugation at 4 °C and 25000 xg for 30 min.

Enzyme assays were performed in 50 mM MOPS buffer (pH 7.8) at 30 °C. Enzyme assays with cell-free extracts contained 0.5 mM Δ^4^-3-keto-UDCA and 0.7 mg ml^-1^ total protein. Activity was determined by HPLC-MS measurement of samples taken at different time points. For all steps of biochemical purification, enzyme activity was tested using enzyme assays with Δ^4^-3-keto-UDCA (V). Substrate concentrations ranged from 0.15 mM in most assays to 0.036 mM in assays after ion exchange chromatography. 0.1 mg ml^-1^ total protein or up to 40 µl of fractions after ion exchange chromatography were used for the assays. Enzyme activity was monitored at 290 nm in a spectrophotometer during incubation at 30 °C for 10 min. Activity of purified Hsh3 and Hsh2 was analyzed in either discontinuous or continuous assays, which were either monitored by HPLC-MS measurements of samples taken at different time points, or monitored spectrophotometrically, respectively. For discontinuous assays, 0.15 mM up to 1 mM substrate as well as 5 µg ml^-1^ up to 200 µg ml^-1^ protein were added. For continuous assays, 50 µM up to 1 mM substrate and 1 µg ml^-1^ up to 5 µg ml^-1^ protein were added and absorption at 300 nm was measured against an assay solution without enzyme. Substrates for these assays were 7α-HADD (XI) and 7β-HADD, which were solved in ethanol. Therefore, ethanol was added to all assays so that the final concentration was always 25%.

### Preparation of steroid compounds

UDCA was purchased from ChemPUR (Karlsruhe, Germany), CDCA from Carl Roth (Karlsruhe, Germany), and cholate from Sigma Aldrich (St. Louis, Mo, USA). Δ^4^-3-keto-UDCA (V) and Δ^4^-3-keto-CDCA (IV) were produced biotechnologically from UDCA and chenodeoxycholate as described previously (24). For this, 1 mM of the respective bile salt was transformed to the Δ^4^-3-keto derivative by strain Chol1 Δ*stdA1* Δ*kstD1* with additional 12 mM succinate until all substrate was transformed. Products were purified from culture supernatants by organic extraction with ethyl acetate after acidification. 7α-hydroxy ADD (7α-HADD, XI) and 7β-hydroxy ADD (7β-HADD) were produced biotechnologically from CDCA and UDCA, respectively. For this, 1 mM of the respective bile salt was transformed with strain Chol1 Δ*kshA* (31). Cultures were incubated until all substrate was transformed and products were purified from culture supernatants by organic extraction with dichloromethane.

For purification of steroid compound P2, cultures of strain Chol11 Δ*hsh3* were grown with 1 mM UDCA until only P2 was found in the supernatant. The compound was extracted by organic extraction with ethylacetate after acidification with HCl. It was resolved in H_2_O_MQ_ and pH was adjusted with NaOH. For testing the stability of P2 under different conditions, the pH in aliquots was adjusted to <2 and >12 using HCl and NaOH, respectively. Further purification of P2 for NMR measurements was achieved by semi-preparative HPLC. It was then extracted from the eluate using organic extraction, resolved in methanol, and dried.

Concentrations of steroid solutions were determined photometrically using the respective absorption coefficients depending on the steroid nucleus configuration (Δ^4^-3-keto and Δ^1,4^-3-keto compounds: 14.7 mol^-1^ cm^-1^, Δ^1,4,6^-3-keto compounds: 13.2 mol^-1^ cm^-1^ (28, 65)).

### Analytical methods

Supernatants of steroid-containing samples were analyzed as described (24, 66) by HPLC-MS measurements after centrifugation for 5 min at >16,000 xg at room temperature.

HPLC-MS measurements were performed using a Dionex Ultimate 3,000 HPLC (ThermoFisher Scientific, Waltham, Massachusetts, United States) with an UV/visible light diode array detector and a coupled ion trap mass spectrometer (Amazon speed, Bruker; Bremen, Germany) with an electro-spray ion source (ESI). Separation was achieved over a reversed phase C18 column (Eurospher II 100-5, 150 × 3 mm, 5 μm particle size; Knauer, Berlin, Germany) at 25 °C. 20 µl sample were injected. Either a method using ammonium-acetate buffer (10 mM, pH 6.7) and acetonitrile at a flow rate of 0.3 ml min^-1^ with a gradient from 10% to 48% acetonitrile (66), or a method using ammonium-acetate buffer (10 mM) with 1% formic acid and acetonitrile at a flow rate of 0.3 ml min^-1^ with a gradient from 10% to 90% acetonitrile (24) was used. For MS detection, the following settings were used: ultra-scan mode, scan range of 50 – 1,000 Da, dry gas flow 12 l min^-1^, dry gas temperature 200 °C, nebulizer gas 22.5 psi, polarity alternating, end plate offset 500 V, and capillary voltage 4,000 V.

Steroid compounds were identified by mass, UV spectrum, and comparison with standards.

### Protein identification

Protein identification was performed with SDS-PAGE as well as native PAGE separated protein. Gel slices were cut into smaller pieces (about 1 mm^2^) prior to washing, reduction, alkylation, and tryptic digestion (67). Separation of peptides was performed with a nano-LC system (Ultiamte 3000 nanoRSLC, Thermo Fisher Scientific, Dreieich, Germany) operated in trap-column (3 µm beads, 75 µm inner diameter, 2 cm length; Thermo Fisher Scientific) mode. Separation was achieved with a 25cm column (75 µm inner diameter, 2 µm beads; Thermo Scientific) applying a linear 90 min gradient from 2% v/v acetonitrile to 50% v/v acetonitrile with subsequent re-equilibration. The nanoLC eluent was continuously analyzed by an online coupled ion-trap mass spectrometer (amaZon speed ETD; Bruker Daltonik GmbH, Bremen, Germany) via an electrospray ion source (captive spray ion source; Bruker Daltonik GmbH) operated in positive mode. Per full scan MS, 20 tandem MS spectra of the most intense masses were acquired (precursor charge 2+ or more) with subsequent active precursor exclusion for 0.2 min. Protein identification was performed with the ProteinScape platform (version 3.1; Bruker Daltonik GmbH) on an in-house Mascot server (version 2.3; Matrix science, London, UK) against a genomic database of *Sphingobium* sp. strain Chol11 translated into amino acids, applying a target decoy strategy, including a mass tolerance of 0.3 Da for MS and 0.4 Da for MS/MS searches and applying a target decoy strategy (false discovery rate <1%) (68).

### NMR analysis

NMR analysis was performed using a Bruker NEO 500 MHz spectrometer equipped with a cryogenically cooled Prodigy HCN-TCI probe. 5 mg of dry sample were dissolved in 280 µL of the deuterated methanol (Deutero GmbH, Kastellaun, Germany) and transferred to a 5 mm Shigemi NMR tube (Shigemi Co. LTD, Tokyo, Japan) matched for methanol. All NMR spectra were acquired at 298 K and all chemical shifts were referenced relative to the tetramethylsilane signal. Structure elucidation was done using 1D ^1^H and ^13^C, as well as 2D ^1^H-^13^C HSQC, ^1^H-^13^C HMBC, 1,1-ADEQUATE, COSY, TOCSY and NOESY experiments. Spectral windows for 1D experiments were 20 ppm for ^1^H and 20 ppm for ^13^C. For 2D experiments ^1^H spectral window was 12 ppm and ^13^C spectral window was 165 ppm for HSQC and 220 ppm for HMBC experiments, respectively. 1D spectra had 65536 data points and 2D spectra had up to 4096 data points in the direct dimension and up to 1024 data point in the indirect dimension. NOESY mixing time was set to 500 ms. All NMR spectra were acquired, processed, and analyzed with Bruker TopSpin v4.1.3 software.

### Bioinformatic methods

Primers for unmarked gene deletion were designed using primerBLAST (69). Hsh3 homologs in other strains and in the database of the human genome project (53) were identified using Hsh3 and BLASTp (63, 70). Putative BaiI homologs in strain Chol11 were also identified using BLASTp. The sequence of Hsh3 was analyzed for domains and protein families using NCBI CDD and Interpro (71, 72). Similarities between Hsh3 and other proteins were calculated using global alignments (70, 73). Hsh3 and Hsh2 homologs in metagenomes of the MGnify database were identified using the MGnify website (74) and phmmer (75). The structure of Hsh3 was predicted using AlphaFold 3 with the standard parameters (76) and visualized from these data using ChimeraX (77).

## Acknowledgements

The authors thank Karin Niermann and Kirsten Heuer for excellent experimental support and Johannes Holert for help with data retrieval. This work was funded by two grants of the Deutsche Forschungsgemeinschaft (DFG projects PH71/3-2 and INST 211/646-1 FUGG) to BP. FMM was supported by a grant of the Deutsche Forschungsgemeinschaft (DFG project number 504745114) during the finalization of the manuscript.

